# Chrysomycin A inhibits the topoisomerase I of *Mycobacterium tuberculosis*

**DOI:** 10.1101/2021.06.23.449690

**Authors:** Balaji Muralikrishnan, Lekshmi K. Edison, Azger Dusthackeer, G R Jijimole, Ranjit Ramachandran, Aravind Madhavan, Ajay Kumar Ramakrishnan

**Affiliations:** Mycobacterium Research Laboratory, Rajiv Gandhi Centre for Biotechnology, Thiruvananthapuram, Kerala, India; Department of Bacteriology, National Institute for Research in Tuberculosis, Chennai, Tamil Nadu, India

## Abstract

Novel anti-tuberculosis drugs are essential to manage drug resistant tuberculosis, caused by the notorious pathogen *Mycobacterium tuberculosis.* We recently reported the antimycobacterial activity of chrysomycin A in vitro and in infected macrophages. In this study, we report that the molecule inhibits the growth of drug resistant clinical strains of *Mycobacterium tuberculosis* and acts in synergy with anti-TB drugs such as ethambutol, ciprofloxacin and novobiocin. In pursuit of its mechanism of action, it was found that chrysomycin A renders bactericidal activity by interacting with DNA at specific sequences and by inhibiting topoisomerase I activity of *Mycobacterium tuberculosis*. It also exhibits weak inhibition of gyrase enzyme of the pathogen.

## Introduction

Chrysomycin A was first discovered by Strelitz et al., in 1955 when screening extracts of actinomycetes against bacteriophages ^1^. Although it was discovered in the golden era of antibiotic discovery, it remained non-available. A few studies have reported its anti-tumorigenicnature ^2–4^. Acetylated forms of chrysomycins were also reported for the same activity ^5^. However, our laboratory was the first one to report chrysomycin A’s anti-*Mycobacterium tuberculosis* property. In our drug screening program against *M. tuberculosis*, we chanced upon chrysomycin A from a novel *Streptomyces* sp ^6^. It has bactericidal activity on both planktonic as well as phagocytosed bacteria. Subsequently in an independent study, derivatives of chrysomycin A have been reported for strong growth inhibitory activity against drug resistant *M. tuberculosis* ^7^. Structurally, chrysomycin A is a napthocoumarin and has a planar structure and this led us to speculate about its interaction with DNA ^8, 9^. Identification of the target of chrysomycin A in *M. tuberculosis* is essential to understand its mechanism of action which ultimately could lead to rational design of potent molecules from its chemical backbone.

This study focuses on identifying the target of chrysomycin A in *M. tuberculosis*. Towards this, we tested the molecule on various drug resistant clinical strains of *M. tuberculosis* to draw initial cues from the drug resistance pattern. Subsequently, DNA interaction studies were carried out using physico-chemical methods and molecular docking analysis. The results were suggestive of inhibition of topoisomerases and to check this possibility, functional inhibition assays were performed and support was drawn from molecular docking analysis.

## Materials and Methods

### Bacterial strains, cell lines, chemicals and general procedures

*M. tuberculosis* H37Rv was grown on Middlebrook 7H9 medium (BD Difco, New Jersey) supplemented with 10% Oleic acid-Albumin-Dextrose-Catalase (OADC, Difco). The details of the drug resistant clinical strains of *M. tuberculosis* are provided in the Table 1. Middlebrook 7H10 solid medium was procured from BD Difco, while flat opti bottom 96 well plates were from ThermoFisher Scientific, Massachusetts, United States. *M. tuberculosis* gyrase and topoisomerase I inhibition kits were bought from Inspiralis (Norwich, UK). Potssium iodide, Sodium chloride and oligonucleotides (listed in supplementary table 2) were procured from Sigma (India)

**Table 1:** Antimicrobial activity of chrysomycin A against drug resistant clinical strains of *M. tuberculosis.*

| Strain* | Resistant to | Susceptible to | MIC(μg/ml) |
| --- | --- | --- | --- |
| 1 | INH, RMP, Km, LFX | No data | 4 |
| 2 | INH, RMP, LFX | Km | 2 |
| 3 | INH, RMP, LFX | Km | 4 |
| 4 | INH, RMP, LFX | Km | 1 |
| 5 | INH, RMP, LFX | Km | 1 |
| 6 | INH, RMP, Km | LFX | 2 |
| 7 | INH, RMP, EMB, M, Ofx | Km, Ak, M | 8 |
| 8 | INH, Ofx | RMP, Km | 16 |
| 9 | INH, RMP, EMB, M, Ofx | Km,Ak, M | 8 |
| 10 | INH | RMP, Km,Ofx | 16 |
| 11 | RMP | INH, EMB,Km,Ak, M, Ofx | 1 |
| 12 | INH, RMP,Ofx | EMB,Km,Ak, M | 16 |
| 13 | RMP, Km | INH, EMB, Ofx, Ak, M | 8 |
| 14 | STR, INH, RMP, EMB, Ofx | Km | 1 |
| 15 | INH, RMP, EMB, Ofx | Km, Ak, M, Ofx | 16 |
| 16 | STR,INH, RMP | EMB,Km,Ofx | 16 |
| 17 | STR,INH, RMP, EMB, | Km, Ofx | 2 |
| 18 | INH | RMP, EMB,Km,Ofx,Ak, M | 4 |
| 19 | INH, RMP,Km, LFX | STR, EMB, Ak | 4 |
| 20 | INH, RMP, Km, | STR, LFX | 2 |
| H37Rv | - |  | 4 |

INH – Isoniazid, RMP – Rifampicin, EMB – Ethambutol, Ofx – Ofloxacin, LFX – Levofloxacin, STR – Streptomycin, Km – Kanamycin, M – Moxifloxacin, Ak – Amikacin.
\* - identity of the clinical isolates withheld.

### Antimicrobial activity of chrysomycin A on drug resistant clinical strains of *M. tuberculosis*

Chrysomycin A was tested on drug resistant clinical strains of *M. tuberculosis*. For this, well characterized mono-drug resistant, multidrug resistant and extensively drug resistant strains were used and micro-dilution assay was performed on each (1-16 μg/mL). After incubating the culture at 37 °C for 24 h, 5 μL of the treated and control cultures were spotted onto Middlebrook 7H10 solid medium. The lowest concentration at which no colonies appeared was considered as the MIC.

### Study of synergism with anti-TB drugs

To check whether chrysomycin A can interact with some of the current anti-TB drugs, checker board assay was performed as described ^10^. Surrogate organism, *Mycobacterium smegmatis,* was used for this particular experiment. First-line drugs, isoniazid, rifampicin and ethambutol were included in the study along with ciprofloxacin and novobiocin. Pyrazinamide, a first-line drug was not included in the study as the drug is not active against *M. smegmatis*. In the checker board assay, synergy between chrysomycin A and the first-line drugs in eliciting antimicrobial property was determined individually. Briefly, as shown in Figure S2A, in a 96 well plate, chrysomycin A was serially diluted (6 μg/mL to 1/16 μg/mL) in horizontal direction from left to right until the penultimate column and leaving the wells in the last column without chrysomycin A. Next, the second line drug (ciprofloxacin, as shown in Figure 32A) is serially diluted (1 μg/mL to 1/16 μg/mL) in the vertical direction from top to bottom until the penultimate row leaving the last row without ciprofloxacin. Thus, the last row and the last column will indicate the individual MIC of chrysomycin A and ciprofloxacin, respectively. The test organism, *M. smegmatis* (1.5 × 10^6^ bacteria) was inoculated and resazurin microtitre assay was performed as described ^11^. The wells between the last row and last column of the 96 well plate bear the different combinations of both the drugs.

### Fluorescence spectrometry

The experiment was performed as described by Rehman et al ^12^ with minor changes. Briefly, flat opti-bottom Nunc 96 well plates were used instead of cuvettes. Chrysomycin A (5 or 50 μM) was kept at constant concentration and its intrinsic fluorescence was read between 400 and 600 nm when excited at 280 nm. Increasing concentrations (0-50μM) of calf thymus DNA or short stretches of single/double stranded DNA were added to this, and the change in fluorescence spectrum was monitored. The spectra were read using JASCON fluorometer and the graph was plotted using Graphpad Prism version 7 Software. Stern-Volmer quenching constant was calculated from the slope of [F/F_0_] vs [DNA] plot. F_0_ is the initial fluorescence before adding potassium iodide and F is the fluorescence obtained after addition, while [DNA] is the concentration of DNA. The constant was compared to that of reported intercalators and groove binders ^13^.

### Effect of potassium iodide and sodium chloride on the fluorescence of DNA-chrysomycinA complex

Potassium iodide quenching was carried out as described by Sadeghi et al ^14^. Briefly, potassium iodide (0–8 mM) was added to chrysomycin A (50 μM) and DNA-chrysomycin complex (10 μM of double stranded DNA with 50 μM of chrysomycin A). The total reaction volume was 200 μL. After exciting at 280 nm, the fluorescence emission spectra were recorded between 400 to 600 nm. A graph was plotted with the ratios of F and F_0_ against increasing concentrations of potassium iodide. The effect of ionic strength was studied by varying the concentration of sodium chloride between 0 and 70 mM in a total volume of 200 μL containing 50 μM chrysomycin A or calf thymus DNA-chrysomycin A complex (1:1). The change in spectrum was recorded at 505 nm and a graph was plotted against increasing concentrations of sodium chloride.

### Circular dichroism (CD)

CD spectra were recorded from 200 nm to320 nm with a scan speed of 200 nm/min with a spectral bandwidth of 10 nm using a JASCON CD spectrophotometer (Maryland, USA). Three scans were averaged to plot the spectra. Concentration of single/double stranded/calf thymus DNA was kept constant at 50 μM and its spectrum was read between 220 and 300 nm. Increasing concentrations of chrysomycin A (0-220μM) were added to DNA and the change in CD spectra was recorded, and the graph was plotted using Graphpad prism version 7 software. The spectra of buffer solution (5mMTris-HCl, pH 8.0) containing chrysomycin A at appropriate concentrations were subtracted from the spectra of DNA and chrysomycin A-DNA complex.

### Interaction of chrysomycin A with single/double stranded oligonucleotides

Antibiotics and drugs with antimicrobial activity that have preferences for DNA sequences were identified from published literature ^15, 16^, and a random set of oligonucleotides was synthesized as short stretches (single stranded) and were used in fluorescence spectrometry as described earlier (section 5.1.1). The most preferred sequences were identified based on the highest fluorescence read out. After identifying these sequences, short oligos of DNA sequences were synthesized and ligated to prepare double stranded DNA, and were used in the fluorescence spectrometry and circular dichroism studies.

### Molecular docking analysis

Protein structures, 5D5H (*M. tuberculosis* topoisomerase I), 5BTD (*M. tuberculosis* gyrase); and DNA structures, IBNA (dodecamer without ACGT site) and IHQ7 (with ACGT site) were retrieved from PDB (protein data bank) database in PDB format. The ligands attached with the protein were removed using PrepWizard software in Schrondinger suites 2019-1. Energy minimization was carried out along with optimizations such as building missing loops and removal of water molecules. SiteMap was used to predict the ligand binding sites and the catalytic domain was selected. Two dimensional structure of chrysomycin A (CID 73468) was retrieved from PubChem database and imported to LigPrep software of the suite and energy minimizations were performed. Finally, with the default setting of 15Å × 15Å × 15Å grid points, a grid was generated. This was used in the docking analysis using Glide dock-XP mode (extra precision).

### Mobility shift assay

To 200 ng of plasmid DNA (pBR322), chrysomycin A was added at different concentrations (0 – 10 μM) and allowed to interact. From each of these reactions, 10 μL of the mix was run on a 1% agarose gel (prepared in 1X Tris acetate buffer, pH 8.0) for 60 min at 50 V/cm in 1X Tris acetate buffer containing ethidium bromide. The DNA band pattern was documented using Bio-Rad gel documentation system.

### Scanning electron microscopy

Fifty microliter of *M. tuberculosis* H37Rv culture (of McFarland standard 1.0 treated with chrysomycin A at 1X MIC overnight, and untreated control) was centrifuged at 8000 g for 10 min at 25 °C. The resulting pellet was washed with ice-cold 1X PBS and suspended in 1.0 mL of Dubos Difco broth. Two hundred microliter of this suspension was loaded onto polylysine-coated cover slips that were prepared according to manufacturer’s protocol (Sigma-Aldrich). The bacteria were allowed to settle on the cover slip for 30 min. Then, the culture was decanted gently and 2.5% glutaraldehyde in PBS (pH 7.4) was added to the coverslip. This was incubated overnight at 4 °C and then subjected to dehydration with a series of ethanol concentrations (50%, 70%, 90% and 100%). At each step of ethanol gradient treatment, 10 min of incubation was provided for effective dehydration. The coverslips were dried at room temperature and the samples were gold-sputtered using JEOL-1200 (Peabody, MA, USA) and imaged using a JEOL-JSM-5600 LV (USA) scanning electron microscope.

### *M. tuberculosis* topoisomerase I inhibition assay

*M. tuberculosis* topoisomerase I enzyme was procured from Inspiralis, UK. Supercoiled pUC19 plasmid was isolated using either a plasmid maxiprep kit (Magerey Nagel) or using a modified method described by Carbone et al. 2012 using miniprep kit (Magerey Nagel). Two units of the enzyme were used in each reaction to relax the supercoiled DNA in the presence of different concentrations of chrysomycinA (10, 20, 40 and 80 μM). Experimental procedures were followed as per the protocol provided by the manufacturer.

### *M. tuberculosis* gyrase inhibition assay

*M. tuberculosis* gyrase enzyme was purchased from Inspiralis, UK. In decatenation assay kinetoplast DNA served as the substrate (every reaction had 200 ng of DNA). Chrysomycin A at different concentrations (5, 50 and 100 μM) was used to inhibit the enzyme function. Supercoiling assay had relaxed topoisomers of pBR322 plasmid (provided in the kit) as substrate, and the assay was carried out in the presence of ATP (1 mM). Chrysomycin A (5, 50 and 100 μM) was used to inhibit the enzyme function. For DNA relaxation assay, pUC19 supercoiled plasmid (300 ng) was used as the substrate and the assay was performed without ATP in the reaction. Ciprofloxacin at 25 μM and novobiocin at 20 μM served as positive control for inhibition. Chrysomycin A at different concentrations (0-80 μM) was used to inhibit the enzyme functions. In all reactions, except in the negative controls, 2 units of the enzyme was used in the decatenation, supercoiling and relaxation assays. All procedures were followed as per the protocol provided by the manufacturer.

## Results and Discussion

### Activity of chrysomycin A against drug resistant clinical strains of *M. tuberculosis*

The antimicrobial activity of chrysomycin A was tested on *M. tuberculosis* strains that were resistant to different drugs and we tested if there is any change in their minimum inhibitory concentrations (MIC). Interestingly, more than 70% of the drug resistant strains recruited were inhibited at the same concentration of chrysomycin A at which it inhibited the susceptible virulent laboratory strain *M. tuberculosis* H37Rv. The remaining strains were inhibited at slightly higher concentrations. The details of the strains, their susceptibility to anti-TB drugs and the MIC observed in the case of chrysomycinA are provided in Table 1.Strains 2-5 were resistant to isoniazid, rifampicin and levoflaxocin while susceptible to kanamycin. However, chrysomycin A elicited bactericidal activity on these strains below its MIC which was observed on *M. tuberculosis* H37Rv. Strain 1 was similar to Strains 2-5 except that it was resistant to kanamycin also. It was killed by chrysomycin A at the same MIC that killed *M. tuberculosis* H37Rv.On the whole, chrysomycin A seemed to be very effective against MDR strains while it was less effective on ofloxacin resistant strains suggesting that the molecule could act on topoisomerases.

### Chrysomycin A exhibits synergistic activity with anti-tuberculosis drugs

To validate the observation on drug resistant strains of *M. tuberculosis*, chrysomycin A was tested along with first-line and second-line anti-TB drugs and checked for their efficacy on bacterial killing. *M. smegmatis* mc^2^155, the surrogate mycobacterium, was treated with chrysomycin A in combination with the first-line drugs rifampicin, isoniazid, ethambutol and second-line drug ciprofloxacin and novobiocin. Figure S2 summarizes the result; briefly, chrysomycin A acts synergistically with ethambutol, ciprofloxacin and novobiocin while it also shows additive property in exerting antimicrobial activity in combination with rifampicin and isoniazid. Chrysomycin A belongs to coumarin class of antibiotics which are known to bind gyrase subunit B and inhibit the enzyme function. Interestingly, while acting in synergy the amount of chrysomycin A and the combination drugs could be reduced by 4 fold to effect the same level of killing. Thus, it seems chrysomycin A could be inhibiting the topoisomerases along with ciprofloxacin and novobiocin. Chrysomycin A showed synergism when treated along with ethambutol, and it could be because the molecule might have secondary targets in the bacterium. Isoniazid and ethambutol are known to act synergistically and are required only at half their original MIC to cause bacterial killing ^17^. However, in combination with chrysomycin A, only one fourth the concentration is required to elicit the same bactericidal activity. Pyrazinamide was not included in the study because it does not inhibit the growth of *M. smegmatis ^18^*.

### Chrysomycin A – DNA interaction

#### Chrysomycin A intercalates DNA at specific sequences

The planar structure of chrysomycin A (Figure S3) and the speculation on its ability to bind DNA warranted DNA interaction studies. The intrinsic fluorescence of chrysomycin A facilitated the drug-DNA interaction studies through the analysis of alterations in the intrinsic fluorescence after forming drug-DNA complex. Chrysomycin A emits fluorescence at 505 nm in TE buffer, (pH 8.0) when excited at 280 nm or 400 nm. Making use of this property, the change in its fluorescence was monitored with addition of increasing concentrations of calf thymus DNA to a constant amount of chrysomycin A. We observed a concentration-dependent increase in fluorescence of chrysomycin A on addition of calf thymus DNA (Figure1A). Also, a red shift (5 nm) was observed in the fluorescence emission after addition of DNA which indicated a complex formation. The shift also shows that chrysomycin A is buried deep inside the hydrophobic DNA helix which prevents its interaction with water molecules. This phenomenon was described by Sirajuddin et al with other intercalating drugs ^19^. The ratio of observed to initial fluorescence was plotted against the respective concentration of DNA and a Stern-Volmer constant (Ksv) of 1.2 × 10^4^ M^−1^ was obtained from the slope of the graph (Figure 1B). The constant obtained matches neither with the lower values of groove binders nor in the higher constants of canonical intercalators (Ksv<10^4^ if it is a groove binder, Ksv>10^5^ in the case of intercalators) demanding a quenching experiment to draw insights into the binding. Therefore to validate the mode of binding, potassium iodide was introduced to quench the fluorescence of unbound and DNA-bound chrysomycin A. Contrary to our expectation, on addition of potassium iodide we observed an increase in fluorescence even from unbound chrysomycin A and the DNA-chrysomycin A complex. A graph was plotted using the F/F_0_ ratios versus the concentration of potassium iodide added (Figure1C). A slope was calculated for both the linear graphs and it was found that the difference in the Ksv (Stern-Volmer constant) was larger indicating that the interaction could be through intercalation. To rule out the possibility of it being a groove binder, increasing concentrations of sodium chloride was added to the DNA and DNA-chrysomycin A complex to increase the ionic strength of the solution which would result in the removal of surface bound molecules and consequently reduce fluorescence intensity. As expected, the fluorescence intensity remained largely unaffected even after the addition of high concentrations of sodium chloride (Figure1D). Further, circular dichroism spectrometry was performed to find chrysomycin A-induced alterations in the secondary structure of DNA. A negative band around 245 nm and positive band around 275 nm are characteristic of B form of DNA (calf thymus DNA). Alteration in the negative band would indicate a change in helicity, and alterations in the positive band would indicate perturbation of the stacks in the DNA ^20^. On treatment with chrysomycin A, the negative band at 245 nm was altered which pointed out at a change in helicity which in turn supported intercalation (Figure 1E). The positive band was disturbed only when very high concentrations of chrysomycin A were added. In addition, to understand whether the molecule could affect supercoiled form of DNA, pUC19 plasmid was treated with chrysomycin A and electrophoresed on an agarose gel. As expected, the supercoiled form of DNA band migrated slowly with increasing concentrations of chrysomycin A (Figure S1). This shift in mobility was also observed by TT Wei et al^9^ who studied interaction of intercalators with plasmid DNA. Furthermore, through molecular docking, chrysomycin A was found to intercalate through minor groove (Figure 1F).

**Figure 1:**
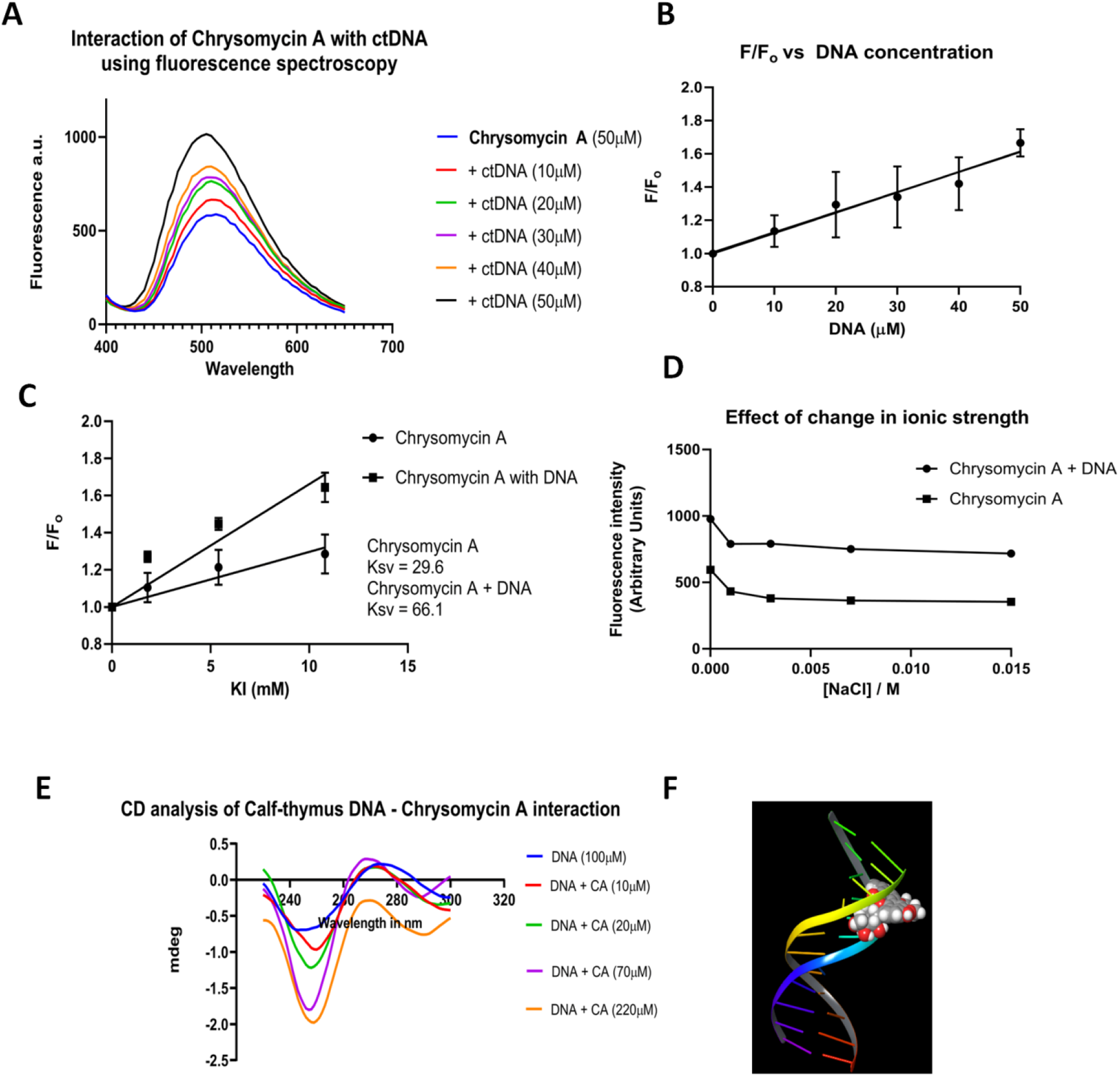
**A**. Change in fluorescence of chrysomycinA on addition of calf thymus DNA. **B.** Determination of Stern-Volmer constant (K_SV_) for DNA-chrysomycin A complex. **C.** Comparison of K_SV_ of free chrysomycin A and DNA-bound chrysomycin A, using potassium iodide as the quencher. **D.** Effect of change in ionic strength on fluorescence of DNA-chrysomycinA complex. **E.** Circular dichroism spectrometry of DNA-chrysomycinA complex. **F.** Molecular docking analysis: chrysomycin A docked onto IBNA.

Therefore, the above evidences confirm that chrysomycin A intercalates DNA with low Stern – Volmer constant (Ksv). This also leads to a hypothesis that the lower affinity observed might be due to chrysomycin A’s preference for specific DNA sequences. By data mining the literature, we could identify such small molecules that bind specific DNA sequences, and they are listed in Supplementary Table 1. Some of those specific sequences were randomly chosen, synthesized and we performed fluorescence spectrometry with chrysomycin A (5 μM).A concentration-dependent increase in fluorescence was observed upon addition of increasing amounts of oligonucleotides, and a significant variation in fluorescence was observed based on the sequence of the oligonucleotides (Figure 2A). This varied fluorescence symbolizes preference for specific sequences. On comparing the sequences, it was found that a GC flanked by A and T appeared to be the most preferred sequence for binding of chrysomycin A. To validate this observation, double stranded DNA oligos with increasing number of AGCT and ACGT sites were synthesized and fluorescence spectroscopy analysis was carried out. As expected, there was an increase in fluorescence depending on the increasing number of AGCT/ACGT sites (Figure 2B and 2D). Change in the secondary structure was also monitored for each of these oligos through circular dichroism spectrometry (Figure 2C and 2E). A significant change was observed in the helicity (increasing negative band at 245 nm) only when there was a single AGCT/ACGT site. Also, oligos containing single ACGT was observed to be more vulnerable to chrysomycin A than those containing AGCT in terms of alterations in the DNA structure. However, this change was not observed in DNA with more than one AGCT/ACGT sites. This is presumed to be due to a stabilization effect of saturated DNA–chrysomycin A complex which does not allow any more change in the secondary structure. To confirm these results, molecular docking was performed with DNA sequences with and without the ACGT sites (1HQ7 and IBNA, respectively) (Figure 3). Docking scores were ten-fold high (−5) in the case of binding to DNA with ACGT site compared to DNA without ACGT site. Also, chrysomycin A intercalated through the major groove in ACGT-containing DNA compared to the minor groove intercalation with DNA lacking ACGT. Hoechst 33238 and ethidium bromide served as controls which have no sequence specificity for binding. Next, in pursuit of finding the significance of the ACGT/AGCT sites, we found an orphan ACGT site in the strong topoisomerase sites (STS) of topoisomerase I of *M. smegmatis* and *M. tuberculosis ^21^*. Interestingly, we found that actinomycin D also exhibits sequence specific binding and binds preferentially to GC flanked by A and T regions of DNA ^16,22^ and, actinomycin D also restricts topoisomerase I activity in tumor cells ^23^. Therefore we wished to test if chrysomycin A also could possibly inhibit topoisomerases of *M. tuberculosis*.

**Figure 2:**
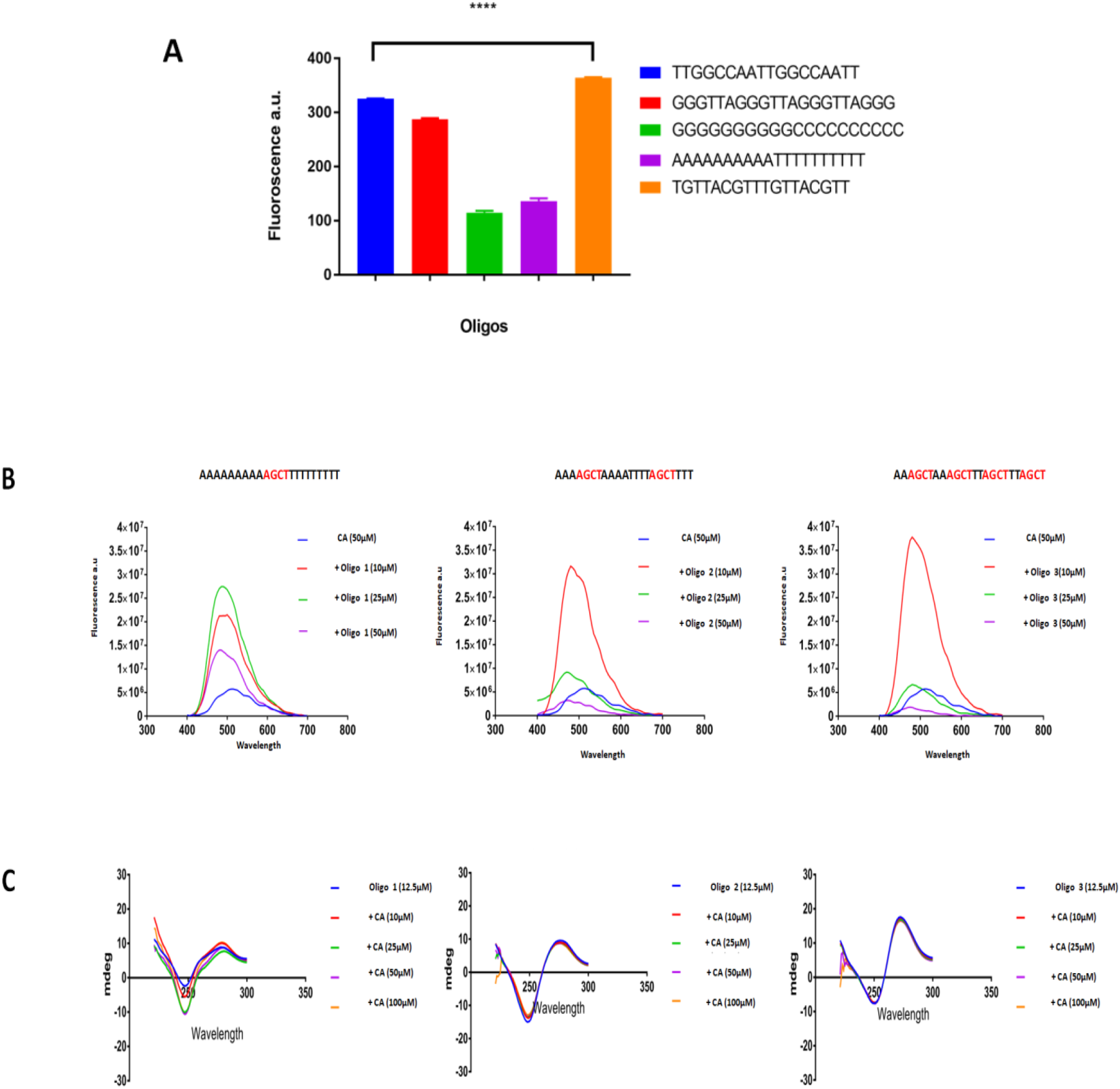
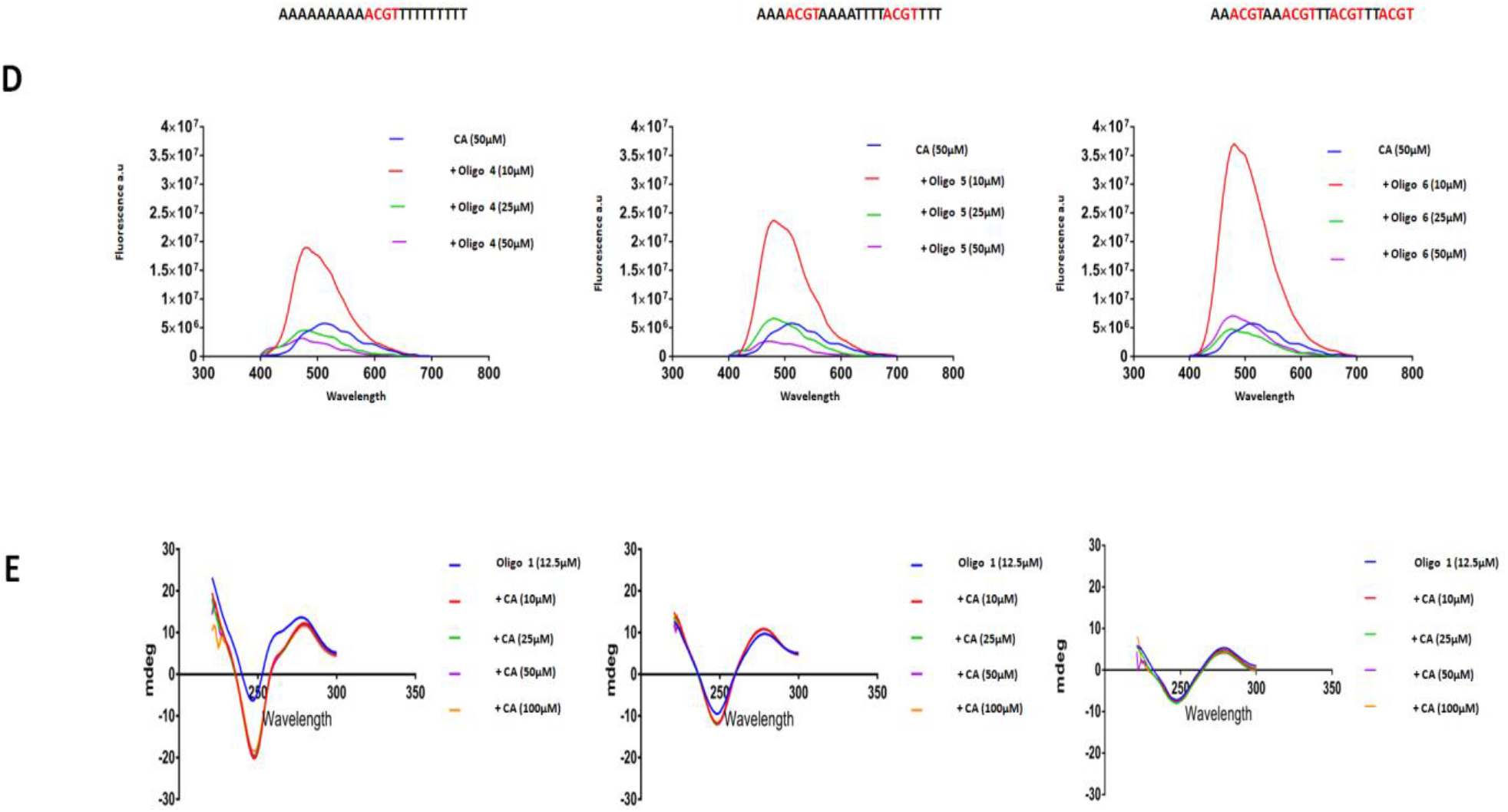
Chrysomycin A preferentially binds to specific nucleotide sequences. A. Fluorescence spectrometry shows differential fluorescence emission of chrysomycinA bound to different nucleotide sequences. B. Change in fluorescence with respect to the number of AGCT sites in the oligos. C. Circular dichroism spectrometry shows changes in the secondary structure of DNA bound to chrysomycin A with respect to the number of AGCT sites in the oligos. D. Change in fluorescence with respect to the number of ACGT sites in the oligos. E. Circular dichroism spectrometry shows changes in the secondary structure of DNA bound to chrysomycin A with respect to the number of ACGT sites in the oligos.

**Figure 3:**
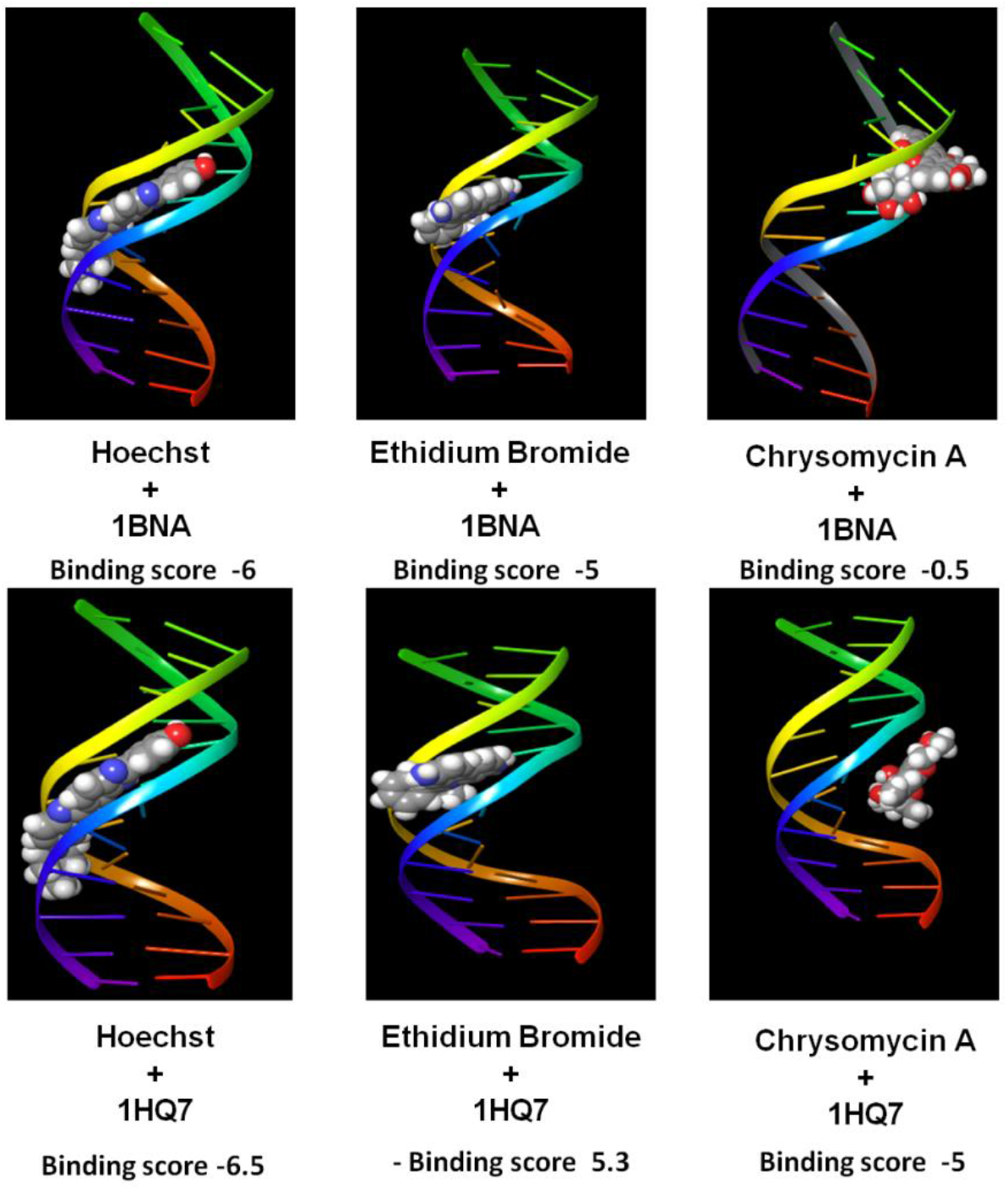
Molecular docking of chrysomycin A, ethidium bromide and Hoechst with DNA sequences with ACGT (1HQ7) and without ACGT (1BNA) site.

#### Chrysomycin A inhibits the activity of topoisomerases of *M. tuberculosis*

Ahmed et al. ^24^ observed that conditional knocking down of topoisomerase I in *M. tuberculosis* resulted in the bacterial shrinkage as well as de-compaction of the genetic materials. A comparable observation was made by Arjomandzadegan et al ^25^, where a drug resistant strain of *M. tuberculosis* treated with ofloxacin (fluoroquinolone that inhibits gyrase enzyme activity) acquired a shrunken abnormal oval shape. Thus, it is evident that inhibition of topoisomerases results in the reduction of bacterial size. To find the effect of chrysomycin A on *M. tuberculosis* morphology, the bacterium was treated with chrysomycin A at its MIC for 12 h and subjected to scanning electron microscopy. We observed 5 to 10 fold reduction in the size of chrysomycin A-treated bacteria when compared to the untreated control (Figure 4A and 4B). We also observed that small rod-shaped bacteria failed to grow further when we tried to retrieve them in an antibiotic-free growth medium. Altogether, the change in morphology hinted at a possible topoisomerase inhibition. In addition, chrysomycin A belongs to coumarin class of antibiotics and coumarins are known for their anti-gyrase activity by binding to the ATP binding subunit B, and therefore we wondered if chrysomycin A might inhibit the gyrase enzyme. Also, we have already shown that it can intercalate at specific sequences in DNA, and intercalators are well known inhibitors of topoisomerase enzymes.

**Figure 4:**
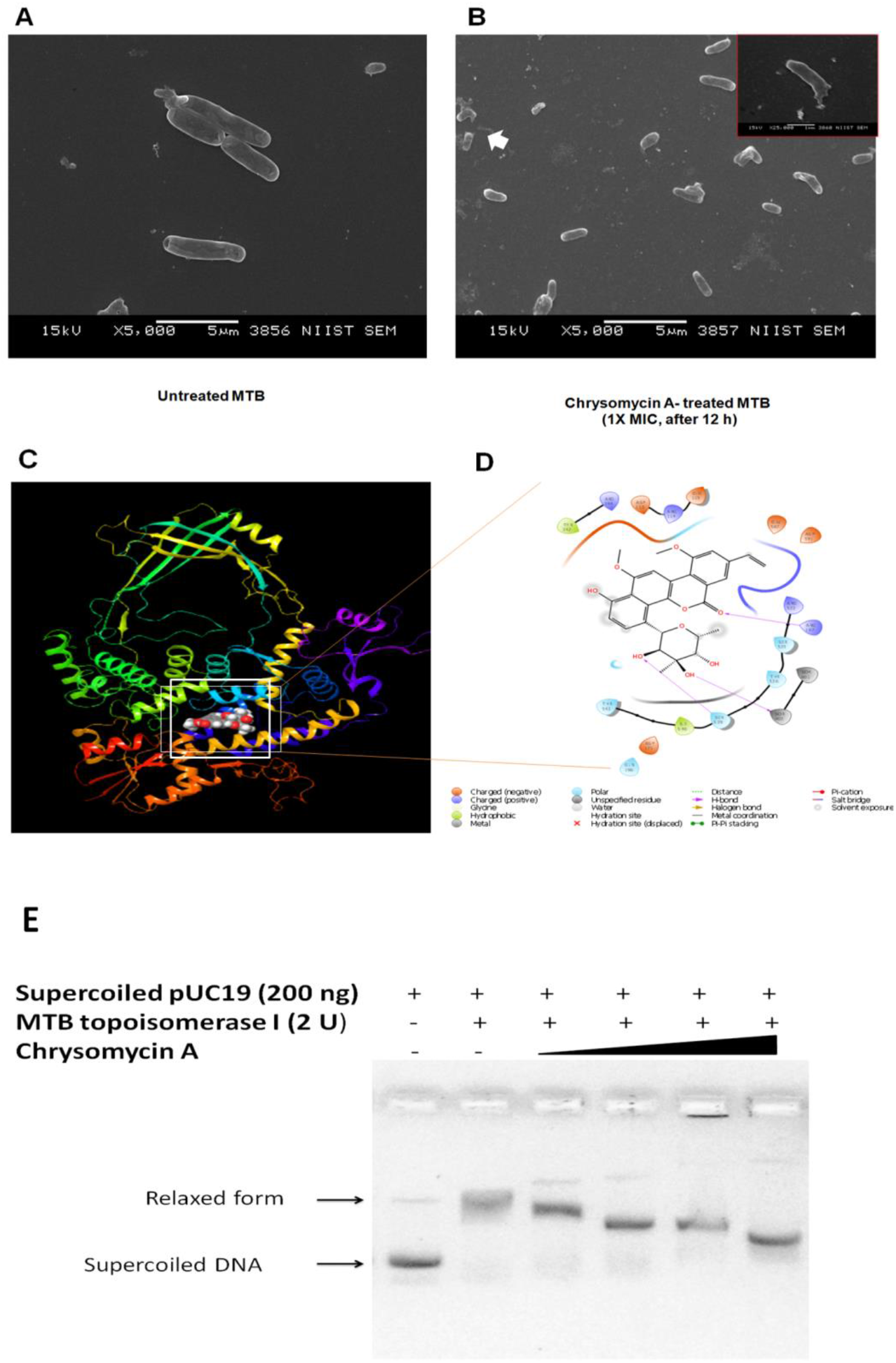
Chrysomycin A interacts with *M. tuberculosis* topoisomerase I to inhibit the growth of the bacteria. A and B are SEM images of untreated and chrysomycin A-treated bacteria, respectively. The inset image in B shows a broken leaky bacterium. C. Molecular docking of chrysomycin A bound to *M. tuberculosis* topoisomerase I protein. D. The binding pocket of chrysomycin A in the protein shows the interacting amino acids. E. DNA relaxation assay with *M. tuberculosis* topoisomerase I and chrysomycin A.

We used Schrondinger docking software to check whether chrysomycin A can bind to *M. tuberculosis* topoisomerase I (PDB ID: 5D5H) and *M. tuberculosis* gyrase (PDB ID: 5BTD). Interestingly, chrysomycin A was found to interact with the primase domain of the topoisomerase I enzyme and the active site Tyr-342 ^26^ was found to interact with the molecule (Figure 4C and 4D). We did not observe any such interaction in the docking study with the gyrase enzyme. Subsequently, a DNA relaxation assay was performed with *M. tuberculosis* topoisomerase I with (0-80 μM) and without chrysomycin A. As expected a concentration-dependent inhibition of relaxation was observed (Figure 4E). At 20 μM, chrysomycin A seemed to inhibit almost half of the enzyme activity. Furthermore, to the best of our knowledge this IC_50_ value of 20 μM seems to be the best among any natural molecule with the same activity. The molecule was also tested for inhibition of various functions of *M. tuberculosis* gyrase. The molecule could inhibit decatenation activity (5 μM) efficiently but exhibited moderate to poor inhibition of the DNA supercoiling (50 μM) and DNA relaxation activity (50 μM) of the gyrase enzyme (Figure 5A, 5B and 5C). We speculate that the gyrase inhibition is through its DNA intercalation rather than through direct binding to the enzyme. Thus, these observations supported our hypothesis that chrysomycin A can inhibit both the topoisomerase I and gyrase enzymes of *M. tuberculosis*.

**Figure 5:**
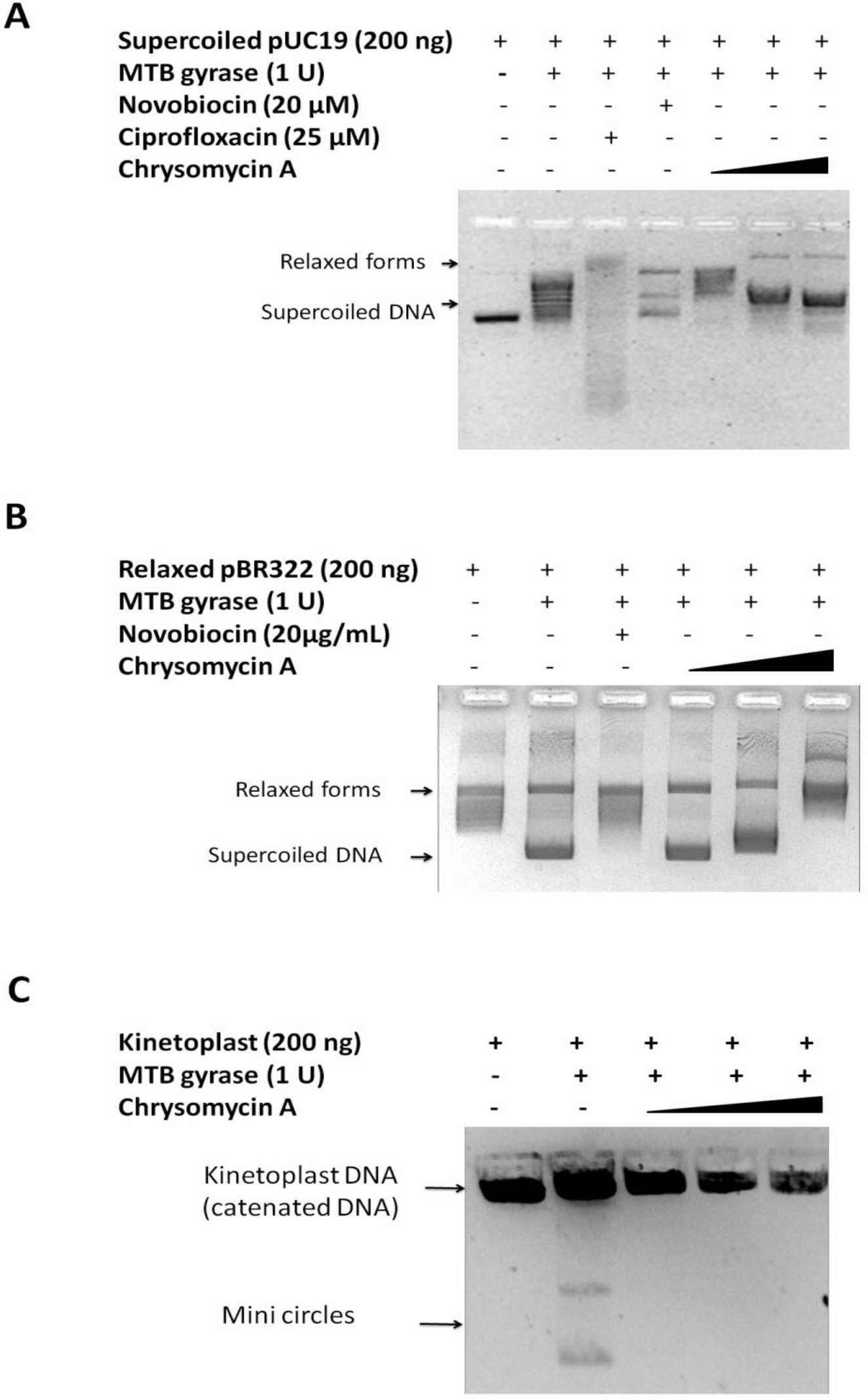
Gyrase inhibition assay. A - DNA relaxation assay. B - DNA supercoiling assay. C - Decatenation assay.

## Conclusion

Chrysomycin A inhibits most of the drug resistant clinical strains of *M. tuberculosis* and acts in synergy with ethambutol, the first line drug and ciprofloxacin, the second line drug. Chrysomycin A inhibits the activity of topoisomerase I and gyrase enzymes of *M. tuberculosis* to kill the bacterium. The inhibition activity was either by binding to the primase domain of the topoisomerase I, or by binding to specific nucleotide sequences of DNA which are apparently the recognition and binding sites of the topoisomerase I, thereby preventing the enzyme activity.

## Acknowledgement

BM thanks Council of Scientific Industrial Research, Govt. of India for research fellowship, and RAK thanks Department of Biotechnology, Govt. of India for funding. We are grateful to Ms. Arthi R and Dr. Kana M Sureshan, School of Chemistry, Indian Institutes of Science Education and Research, Thiruvananthapuram, for their help in fluorescence spectrometry and circular dichroism studies.

